# Mitochondrial Responses to Conventional and Ultra-high Dose Rate (FLASH) Radiation

**DOI:** 10.1101/2025.04.03.647049

**Authors:** Emily G. Caggiano, Alan Lopez Hernandez, Trey Waldrop, Kevin Liu, Henry Gatica-Gutierrez, Sofía Vargas-Hernández, Nefetiti Mims, Ariana Acevedo-Diaz, Brett Velasquez, Denae Neil, Edgardo Aguilar, Matthew D. Meyer, Gloria V. Echeverria, Albert C. Koong, Michael T. Spiotto, Anna-Karin Gustavsson, Emil Schüler

## Abstract

**Purpose:** Ultra-high dose rate (>40 Gy/s, FLASH) radiation therapy (RT) provides equivalent tumor control while reducing normal tissue toxicity relative to conventional dose rate (CONV) RT. However, the mechanisms underlying the observed FLASH effect are unknown. We hypothesized that the preservation of mitochondrial integrity in nontumorigenic cells by FLASH RT could be a key factor in reducing normal tissue toxicity and improving overall treatment outcomes.

**Methods:** We examined mitochondrial health and function after CONV and FLASH in vitro, ex vivo, and in vivo through assays of metabolic flux, mitochondrial membrane potential, mitochondrial reactive oxygen species (ROS), mitochondrial DNA damage and copy number, mitochondrial morphology, and tumor growth and survival.

**Results:** In in vitro assays, murine pancreatic cancer (PDAC) cells showed evidence of equal mitochondrial damage in response to CONV and FLASH, but nontumorigenic pancreatic cells were spared by FLASH. These results were recapitulated ex vivo, and mice treated with FLASH showed higher response rates and longer survival time than mice treated with CONV in an in vivo tumor model.

**Conclusions:** Collectively, these results suggest that FLASH spares mitochondrial function in nontumorigenic cells, but not in PDAC cells, relative to CONV. The preservation of mitochondrial integrity in nontumorigenic cells may be a key mechanism underlying the reduced normal tissue toxicity observed with FLASH RT.

## Introduction

Radiation delivered at ultra-high dose rates (>40 Gy/s, FLASH) can provide equivalent tumor control but less normal tissue damage relative to conventional dose rate radiation (CONV, 0.01-0.40Gy/s), a phenomenon known as the FLASH effect^1–3^. FLASH radiation therapy (RT) is a unique therapeutic modality that may allow delivery of higher-intensity treatment without inducing commensurate toxicity in surrounding tissue^4,5^. However, the mechanisms underlying the observed FLASH effect are unknown^6^.

CONV causes extensive damage to mitochondria, disrupting oxidative phosphorylation, inducing reactive oxygen species (ROS), and causing oxidative stress^7^. In comparison, the effects of FLASH on mitochondrial health and function are understudied and accordingly less well known^3,8,9^. One proposed mechanism for normal tissue sparing afforded by FLASH is related to the generation of ROS and its metabolic pathways^10^. Because mitochondria are the major producers of ROS in cells, differences in mitochondrial biology after FLASH (relative to CONV) may underlie the FLASH effect, but this possibility has not been explored.

One tumor type that would benefit from a better understanding of FLASH mechanisms is pancreatic ductal adenocarcinoma (PDAC), which grows rapidly, metastasizes early, and is lethal in nearly every case. No targeted therapies exist for PDAC, and immunotherapy has limited efficacy owing to its “cold” immune environment^11^. At present, the treatment options available are surgery, for which few patients are eligible given the rapid spread of the disease, or chemotherapy, which is of limited benefit as a monotherapy and often negatively affects quality of life^12–14^.

Although RT is an acceptable treatment option for localized PDAC^15,16^, the radiation dose is often limited by the proximity of the pancreas to radiosensitive organs like the stomach and intestines, which are particularly susceptible to radiation-induced injury such as ulceration and perforation^6^. Thus, additional, more effective treatment options are urgently needed for PDAC patients. FLASH is particularly relevant in PDAC, where curative doses are seldom achieved. FLASH significantly reduces radiation-induced gastrointestinal (GI) toxicity^6,17–20^, thus potentially allowing escalation of the prescribed dose, resulting in improved treatment outcomes.

Here, we endeavored to identify the effects of FLASH, in comparison with CONV, on mitochondrial biology in both PDAC and nontumorigenic ductal pancreatic cell lines. We sought to corroborate our in vitro results with ex vivo and in vivo experiments to evaluate mitochondrial health after both forms of RT. We hypothesized that although CONV and FLASH would both have equivalent effects on PDAC cell lines, FLASH would induce less mitochondrial injury and stress in nontumorigenic cells, resulting in improved mitochondrial function and overall treatment outcome.

## Materials and Methods

### Cell lines and reagents

Murine epithelial *LSL-**K**RAS*^G12D^; *Tr**p**53*^fl/+^; *Ptf1⍺ **C**re* (KPC) cells syngeneic to C57BL/6 mice and squamous *LSL-KRAS*^G12D^; *Trp53*^fl/+^; *Ptf1⍺ Cre* (HY24160) cells^21^ were grown in RPMI-1640 with 10% fetal bovine serum (FBS), 1% penicillin/streptomycin, 2mM GlutaMAX, 1mM sodium pyruvate, and prophylactic plasmocin. KPC media was also supplemented with 7μg/mL insulin. Murine *Trp53*^fl/fl^; *Ptf1⍺ Cre* (IK3025) pancreatic ductal cells^21^ were grown in DMEM/F12 with 15mM HEPES supplemented with 5mg/mL D-(+)-glucose, 1.22mg/mL nicotinamide, 5nM 3,3’,5-triiodo-L-thyronine, 1μM dexamethasone, 100ng/mL cholera toxin, 5ml/L insulin-transferrin-selenium, 1% penicillin/streptomycin, 100μg/mL soybean trypsin inhibitor, 20ng/mL mouse epidermal growth factor, 5% Nu-Serum IV culture supplement, and 25μg/mL bovine pituitary extract. All cells were cultured in a humidified incubator at 37°C with 5% CO_2_, authenticated by short tandem repeat profiling, and tested bimonthly for mycoplasma contamination. Reagents are shown in Table S1.

### Irradiation setup

All irradiations were done with a Mobetron system (IntraOp Medical, Sunnyvale, CA) to deliver 9MeV electrons at CONV and FLASH dose rates^22^. For cell studies, cells were plated in 6cm dishes and irradiated with 4Gy CONV or FLASH (D_50%_ dose). For in vivo studies, tumor-bearing mice were positioned to minimize normal tissue irradiation, and the radiation field was aligned to cover the subcutaneous tumor with a 3mm margin. Mice were irradiated with 20 or 25Gy CONV or FLASH.

Dosimetric calibration for each irradiation was done with dose-rate-independent Gafchromic EBT3 film paired with ion chamber and beam current transformers (BCTs)^23–26^. During FLASH irradiation, the BCTs monitored dose delivery and logged all radiation beam parameters on a per-pulse basis for each delivery.

For animal irradiations, a custom-made collimator was created and positioned at the exit window of the Mobetron as previously described to facilitate sedation and ensure reproducible setup^23,27–29^. The irradiation setup was calibrated for each condition to a depth of 2mm (approximately the center of the tumor) in solid water. Beam parameters are listed in Table S2^30,31^.

### In vivo studies

C57BL/6J mice (8-10 weeks old at the start of all studies) were subcutaneously injected with 1x10^6^ KPC cells and palpated 3 times a week for tumors. When tumors exceeded 100mm^3^ as measured with calipers, mice were irradiated as described above. Tumor volumes were measured 3 times a week until endpoint. The unirradiated controls were placed under anesthesia for the typical treatment time (2-3 minutes) but was not removed from the isoflurane chamber. All mouse procedures used were approved by the Institutional Animal Care and Use Committee (IACUC) at our institution.

### Mitochondrial membrane potential

10mM stock of tetramethylrhodamine, ethyl ester (TMRE) perchlorate was diluted to 100nM working concentration. Cells were stained with TMRE for 30 minutes in a 37°C incubator, protected from light. Unstained control was incubated with media only. After washing, cells were trypsinized and prepared for flow cytometry analysis (AttuneNxT, Thermo Fisher Scientific).

### Oxygen consumption rate assay

OCR was measured as described previously^32^. Briefly, the day before the assay, the XFe24 sensor cartridge was hydrated with calibrant and left overnight in a 37°C non-CO_2_ incubator and cells were seeded at 8,000–15,000 cells per well (optimized per cell line). On the day of analysis, cells were washed and equilibrated for 1 hour in a 37°C non-CO_2_ incubator in XF base assay medium supplemented with 1mM pyruvate, 2mM glutamine, and 10mM glucose. The cell plate was loaded into the bioanalyzer and treated with oligomycin (1.5μM), FCCP (2.5μM), and rotenone A/antimycin (0.5μM) for analysis.

For ex vivo experiments, 50μL of 22.4μg/mL CellTak was added to each well and incubated for 20 minutes. Cells (80,000 per well) were plated, and the plate was centrifuged and incubated at 37°C for 25 minutes to allow cell attachment. 400μL of media was then added to each well, and plates were incubated at 37°C. The assay was then run as described above. All OCR data were normalized to total cell number plated per well.

### Mitochondrial ROS measurement

MitoSOX was dissolved at 5mM stock, then diluted to 5μM working concentration in HBSS. Cells were washed with HBSS and stained with MitoSOX working solution for 15 minutes at 37°C, protected from light. After washing, cells were trypsinized and prepared for flow cytometry analysis (AttuneNxT).

### Mitochondrial DNA damage

The mitochondrial DNA (mtDNA) damage assay was described previously^33^. Briefly, genomic DNA was isolated and quantified. Long-amplification PCR was done in a 25μL reaction containing 2.5 units of Hot Start LongAmp DNA Polymerase, 1x LongAmp Taq Reaction Buffer, 300μM of dNTPs, 2mM MgSO_4_, 0.4μM of each primer, and 50ng of DNA template. PCR product was diluted 1:500 for quantitative PCR (qPCR). qPCR amplifications were done in a 10μL reaction containing 2μL of the 1:500 diluted PCR DNA template, 0.5μM of each primer, and 1x iTaq Universal SYBR Green Supermix. A standard curve was derived from serial dilutions of genomic DNA template to 2, 0.2, 0.02, 0.002, 0.0002, and 0.00002ng DNA. The Ct values were plotted against the logarithm of the dilution, and long-amplification DNA content calculated by linear regression. High DNA content indicates relatively less damaged mitochondrial DNA. Primers are listed in Table S3, thermocycler settings are listed in Table S4.

### Nuclear DNA damage

Cells were fixed with 4% paraformaldehyde, quenched with 10 mM ammonium chloride, permeabilized with 0.2% Triton-X, and blocked with 3% bovine serum albumin. Cells were then immunolabeled for γH2A.x (1:500 dilution) with Alexa Fluor 647-conjugated secondary antibodies (1:1000 dilution) and nuclear stain DAPI. Confocal microscopy was used to acquire images. To quantify γH2A.x foci per nucleus, manual nuclear masks were made from the DAPI channel. Custom MATLAB code was used to process the images and extract γH2A.x cluster counts per nucleus. Extended methods in Supplemental Materials.

### Mitochondrial DNA copy number

Genomic DNA was isolated and quantified. qPCR was done in a 10μL reaction containing 10ng of genomic DNA, 1x iTaq Universal SYBR Green Supermix, and 0.5μM of each primer against murine mitochondrial ND1 and ND6, and nuclear 18S. Raw copy number was calculated using the 2^-ΔCt^ method for each mitochondrial and nuclear DNA primer pair, then averaged together (Table S3 and S4).

### Mitochondrial morphology

Cells were stained with 100nM MitoTracker Orange CMTMRos for 45 minutes at 37°C, protected from light, fixed with 4% paraformaldehyde for 15 minutes at room temperature, and stained with Hoechst (1:3000) for 10 minutes at room temperature. Stained cells were imaged with Airyscan Z-stack confocal microscopy at 63x oil immersion. Morphology was quantified with the ImageJ Mitochondrial Analyzer plugin^34^ to generate a mitochondrial skeleton, from which branches per mitochondria were quantified. Threshold settings were optimized per cell line and treatment condition. Extended methods in Supplemental Materials.

### Western blotting

Cells were lysed with M-PER and protein was quantified with a Bradford assay. Cell lysates were resolved on 4-20% gradient gels and transferred onto PVDF membranes. Membranes were blocked for 1 hour in 5% nonfat milk/TBST. Primary antibodies were diluted 1:1000 in SuperBlock Blocking Buffer and membranes were incubated overnight at 4°C. Secondary antibody was diluted 1:10,000 in 5% nonfat milk/TBST and membranes were incubated for 1 hour at room temperature. For imaging, clarity western ECL substrate was mixed per the manufacturer’s instruction, and blots were imaged with chemiluminescence. Antibodies are listed in Table S5.

### Transmission electron microscopy

Samples were fixed in Karnovsky’s fixative, post-fixed in 1% osmium tetroxide, dehydrated in a graded series of ethanol of increasing concentrations, infiltrated with and embedded in epoxy resin, and heat-polymerized. A Leica EM UC7 ultramicrotome was used to cut 100nm sections, which were stained with uranyl acetate and lead citrate and examined with a JEOL JEM-1400 Flash TEM operating at 120 kV of accelerating voltage and equipped with a high-contrast pole piece and an AMT NanoSprint15 Mk-II sCMOS camera. Mitochondria length was measured in ImageJ.

### Tissue digestion

3.5 days after irradiation, mice were euthanized and tissue was extracted, minced into small pieces, and digested in 1mL of 2x buffer containing collagenase A, hyaluronidase, and bovine serum albumin. Samples were placed in a 37°C shaking incubator for 1-3 hours and vortexed every 15 minutes to aid digestion. Cells were spun down and resuspended in 500μL of ACK red lysis cell buffer for 3 minutes. ACK was inactivated with RPMI media containing 5% FBS and 1x penicillin/streptomycin (epithelial media), and cells were resuspended in 500μL TrypLE for 3 minutes. TrypLE was inactivated with epithelial media, and cells resuspended in 500μL dispase supplemented with 10μL of 1mg/mL DNase1 solution for 3 minutes. Cells were spun down, resuspended in 1mL of epithelial media and filtered through 70μm cell strainers, followed by 40μm cell strainers. Cells were then assessed for viability and counted for downstream assays.

### Statistical analysis

Data are presented as mean ± SEM and normalized to unirradiated controls unless where otherwise stated. Differences between groups were assessed with two-tailed unpaired *t*-tests or ordinary one-way ANOVA using GraphPad Prism. Kaplan-Meier curves were generated with GraphPad, and curves were compared with the log-rank Mantel-Cox test. For each group, at least three independent experiments were performed. ns, not significant; \**P*<0.05; \*\**P*<0.01; \*\*\**P*<0.001; \*\*\*\**P*<0.0001.

## Results

### Mitochondrial function in PDAC cells is affected similarly after CONV and FLASH

To test mitochondrial function after RT, we evaluated mitochondrial membrane potential and OCR. In KPC cells, both CONV and FLASH significantly reduced mitochondrial membrane potential relative to unirradiated 72 hours after treatment (Fig. 1A). In contrast, in HY24160 cells, mitochondrial membrane potential was increased relative to control (Fig. 1B). Neither cell line showed a difference in response between CONV and FLASH.

**Fig. 1.**
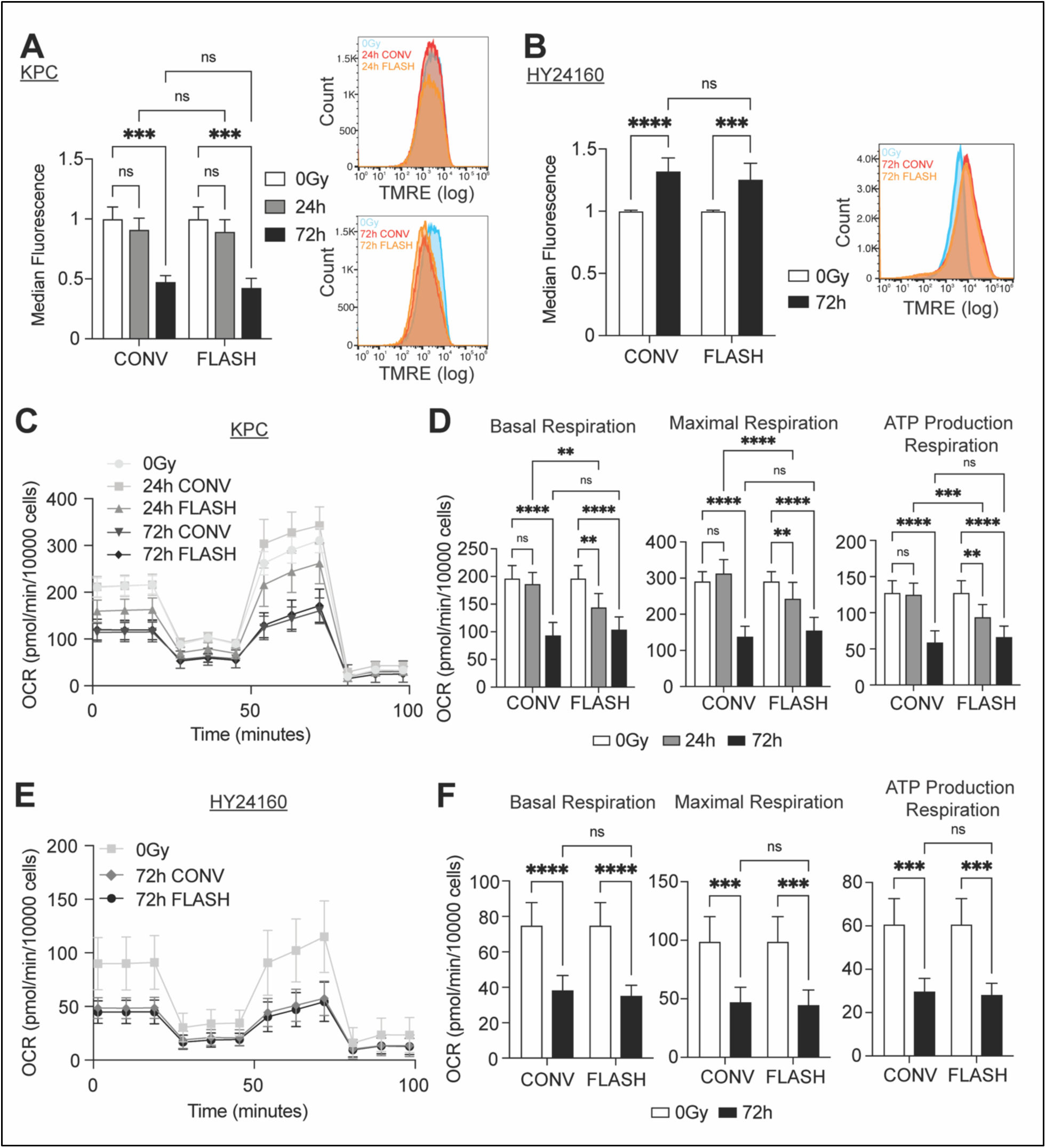
Mitochondrial function in pancreatic ductal adenocarcinoma (PDAC) cells after conventional (CONV) and ultra-high dose rate (FLASH) radiation. Membrane potential in (A) KPC cells 24h and 72h and (B) HY24160 cells 72h after irradiation. (C) OCR in KPC cells 24h and 72h after irradiation with (D) quantification of basal respiration (left), maximal respiration (middle), and ATP-linked respiration (right). (E, F) Corresponding data for HY24160 cells 72h after irradiation.

Next, we evaluated OCR. In KPC cells, CONV and FLASH significantly reduced OCR at 72 hours (Fig. 1C), affecting basal, maximal, and ATP-linked respiration (Fig. 1D). This was reiterated in HY24160 cells (Fig. 1E-F). These findings suggest that although both CONV and FLASH affect mitochondrial function, FLASH does not spare mitochondrial function over CONV in PDAC cell lines.

### FLASH and CONV induce similar mitochondrial stress in PDAC cells

To test mitochondrial stress after RT, we evaluated mitochondrial superoxide generation, mtDNA damage, and mtDNA copy number. In KPC cells, mitochondrial superoxide levels were significantly increased relative to unirradiated controls (Fig. 2A) by 6 hours after irradiation, with no difference in response between CONV and FLASH. Significant mtDNA damage was observed after 6 hours (Fig. 2C), as demonstrated by the failure of the PCR fragment to amplify in the reaction owing to damage-induced DNA fragmentation. Interestingly, at 7 days (168 hours) after RT, CONV-irradiated cells seem to have repaired mtDNA, whereas FLASH-irradiated cells showed a plateau around 70% repaired relative to unirradiated controls. At most time points over all three assays, CONV and FLASH produced similar effects. However, differences between modalities were seen in mtDNA damage 72 hours after treatment and in mtDNA copy number 7 days after treatment. Finally, we evaluated nuclear DNA (nDNA) damage 24 hours after irradiation and determined that CONV had significantly more nDNA damage compared to FLASH (Fig. 2D).

**Fig. 2.**
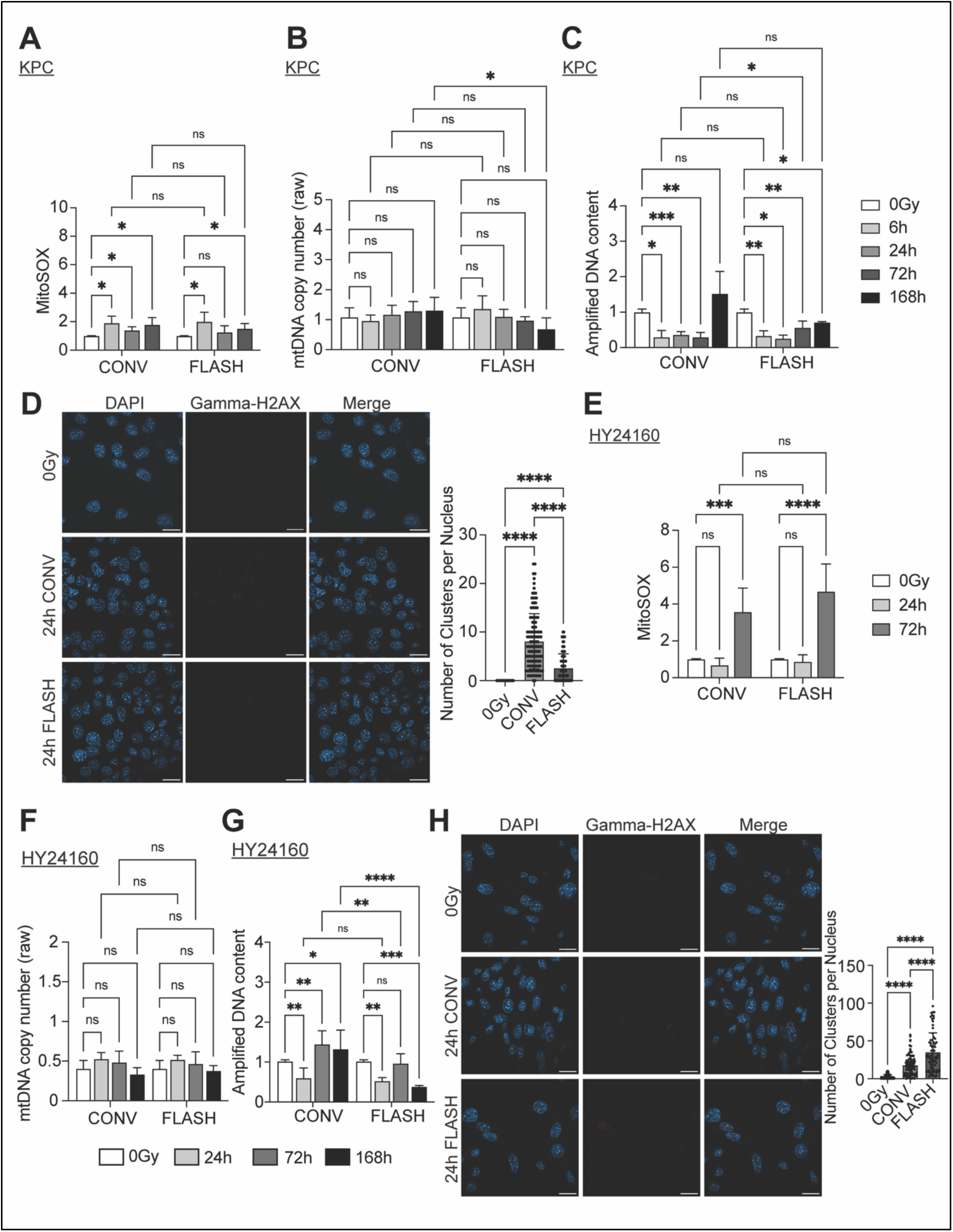
Mitochondrial stress in PDAC cells after CONV and FLASH radiation. (A) Mitochondrial superoxide, (B) mitochondrial DNA (mtDNA) copy numbers, and (C) mtDNA damage levels in KPC cells after irradiation. (D) Nuclear DNA damage levels with γH2A.x staining in KPC cells 24h after irradiation (scale bar 20μm), quantified as number of γH2A.x clusters per nucleus (right). (E-H) Corresponding data for HY24160 cells 24, 72, and 168h after irradiation.

Mitochondrial superoxide levels in HY24160 cells showed similar trends to KPC cells, with significant increases after both CONV and FLASH but no difference between the two modalities (Fig. 2E). mtDNA copy number were also similar, with no difference in treated versus unirradiated conditions and in CONV versus FLASH (Fig. 2F). FLASH induced similar mtDNA damage in HY24160 compared to KPC, with damage remaining after 168h (Fig. 2G), while damage after CONV was reversed. In contrast, FLASH had significantly more damage than CONV 72 and 168 hours after treatment in HY24160 cells. nDNA damage was higher in FLASH compared to CONV 24 hours after radiation (Fig. 2H). These findings indicate that FLASH does not spare mitochondrial stress over CONV in PDAC cell lines with regard to superoxide generation and mtDNA copy number, but the mtDNA damage repair response may differ between the two conditions.

### FLASH induces less mitochondrial stress with reduced impact on mitochondrial function in nontumorigenic pancreatic cells compared to CONV

To evaluate potential differential effects between tumor and nontumorigenic cells after CONV and FLASH, we evaluated mitochondrial stress and function in the nontumorigenic murine epithelial pancreatic ductal cell line IK3025. Mitochondrial membrane potential was unchanged after FLASH but was significantly increased 72 hours after CONV relative to unirradiated controls (Fig 3A). Conversely, mitochondrial superoxide levels were unchanged at all time points tested after CONV but increased significantly 72 hours after FLASH. However, no significant differences were noted between CONV and FLASH at either time point for either assay (Fig. 3A&B). For mtDNA copy number, a significant increase in copy number 72 and 168 hours after CONV was found, but FLASH increased copy number only at 24 hours after irradiation. mtDNA copy number was significantly different 7 days after CONV versus FLASH (Fig. 3C). Regarding mtDNA damage, CONV led to significant damage in mtDNA by 24 hours with recovery by 7 days after irradiation. In comparison, FLASH did not result in any significant mtDNA damage at any time point. However, no significant differences were found between FLASH and CONV (Fig. 3D). Similar to mtDNA, nDNA damage was significantly higher 24 hours after CONV versus FLASH (Fig. 3E). Finally, our evaluation of OCR in IK3025 cells showed that CONV led to significant reductions in OCR, basal, maximal, and ATP-linked respiration at 24 and 72 hours after treatment. FLASH did not induce significant differences in basal and ATP-linked respiration at any time point but showed a significant difference in maximal respiration 72 hours after irradiation (Fig. 3F, G). These findings demonstrate that FLASH largely spares nontumorigenic cells from the mitochondrial stress and loss of function induced by CONV.

**Fig. 3.**
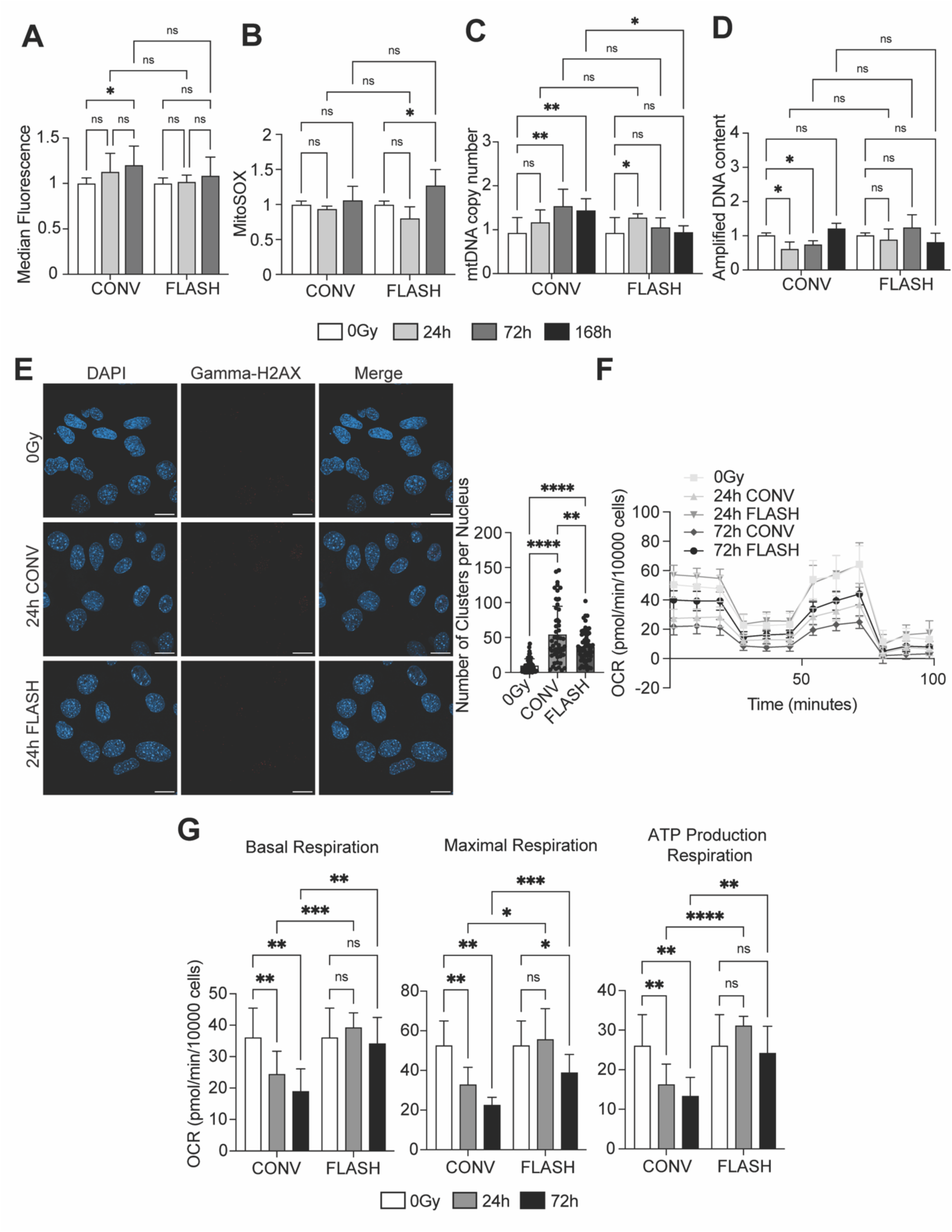
Mitochondrial function and stress in nontumorigenic pancreatic cells after CONV and FLASH. (A) Mitochondrial membrane potential and (B) mitochondrial superoxide levels in IK3025 cells 24h and 72h after irradiation. (C) mtDNA copy number and (D) mtDNA damage in IK3025 cells 24, 72, and 168h after irradiation. (E) Nuclear DNA damage levels with γH2A.x staining 24h after irradiation (scale bar 20μm), quantified as number of γH2A.x clusters per nucleus (right). (F) OCR in IK3025 cells 24h and 72h after irradiation with (G) quantification of basal respiration (left), maximal respiration (middle) and ATP-linked respiration (right).

### Mitochondrial morphology is unchanged after CONV or FLASH in KPC and nontumorigenic cells

We next evaluated the number of branches per mitochondria as a proxy of morphology in PDAC and nontumorigenic cells after CONV or FLASH. In KPC cells, morphology was unchanged up to 72 hours after CONV or FLASH (Fig. 4A). Notably, HY24160 cells had significantly altered mitochondrial morphology compared to the unirradiated control and between CONV and FLASH irradiation (Fig. 4B), as defined by branches per mitochondria. In the nontumorigenic IK3025 cells, morphology was unchanged up to 72 hours (Fig. 4C). In KPC cells, expression of mitochondrial fission and fusion proteins (DRP1, MFN1, MFN2, OPA1) reiterated the morphologic results, with no substantial changes to any protein probed up to 72 hours after CONV or FLASH (Fig. 4D). Moreover, no change was found in either the activating (616) or inhibiting (637) phosphorylating marks on the main mitochondrial fission protein, DRP1, nor the proteins responsible for adding the phosphorylation marks (p/Erk for 616 and PKA for 637). Collectively, these findings demonstrate that mitochondrial morphology is unchanged after CONV and FLASH in KPC and nontumorigenic cell lines at the time points evaluated.

**Fig. 4.**
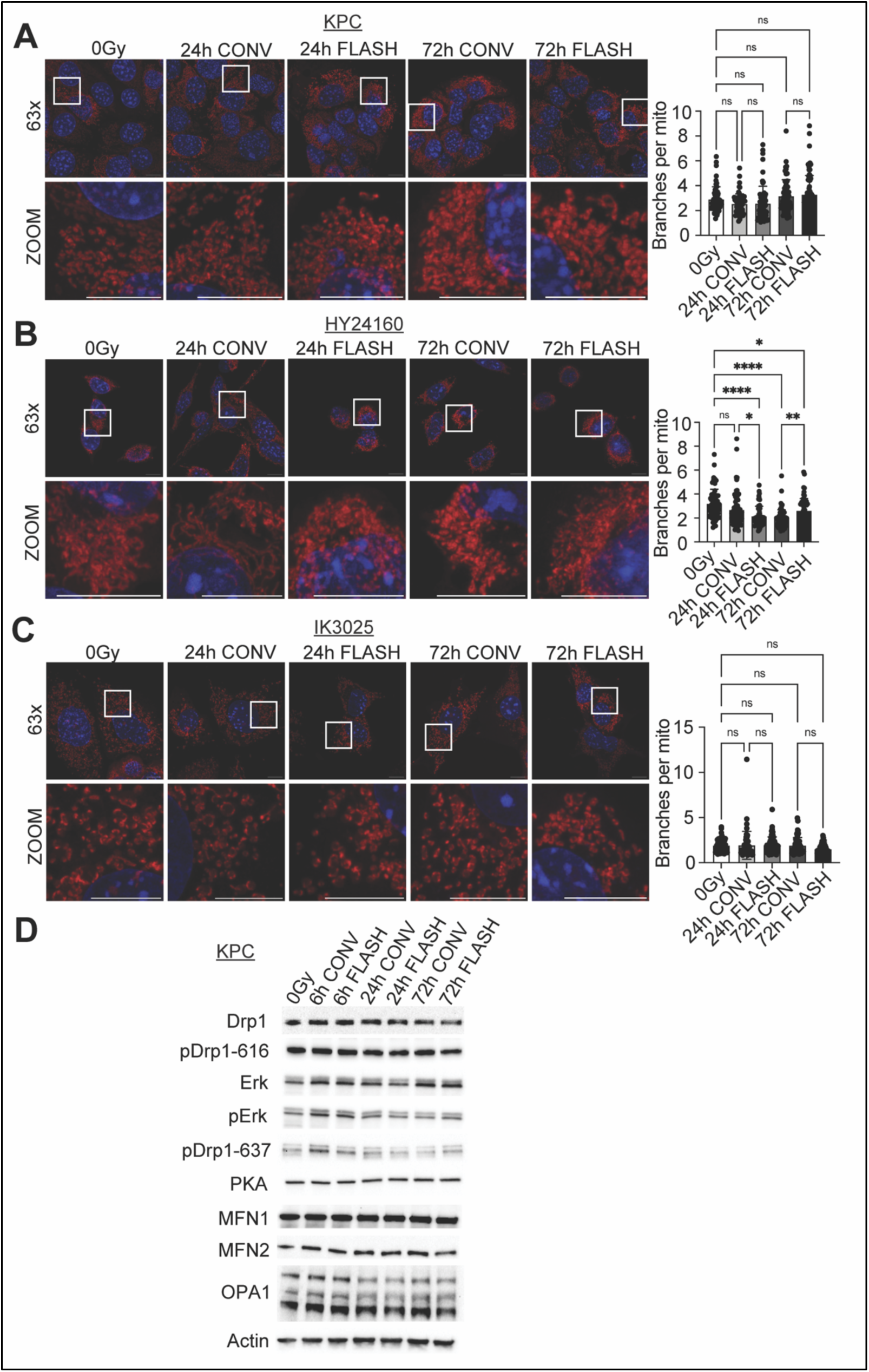
Mitochondrial morphology after CONV and FLASH in pancreatic cancer and nontumorigenic cell lines. Mitochondrial morphology in (A) KPC, (B) HY24160, and (C) IK3025 cells 24h and 72h after irradiation (scale bars 10μm), quantified as branches per mitochondrion (right). (D) Mitochondrial dynamics protein expression in KPC cells 6, 24, and 72h after irradiation.

### Ex vivo measurements reiterate in vitro mitochondrial effects after CONV and FLASH

We next evaluated if KPC tumor cells injected subcutaneously into C57BL/6 mice treated with CONV or FLASH would recapitulate our in vitro results. Evaluation of OCR in tumor cells after excision and digestion revealed significant decreases in OCR, basal, maximal and ATP-linked respiration after 20Gy of either CONV or FLASH (Fig. 5A-B). However, no difference was found between the two modalities. Transmission electron microscopy revealed that although CONV did not alter mitochondrial length in tumors relative to the unirradiated control, FLASH slightly increased mitochondrial length (Fig. 5C). However, in normal pancreas tissue, FLASH did not alter mitochondrial length compared to unirradiated control, but CONV resulted in significantly shorter and rounder mitochondria, with loss of cristae organization (Fig. 5D). These findings demonstrate that the impacts on PDAC mitochondrial function seen in vitro are reiterated in an ex vivo setting.

**Fig. 5.**
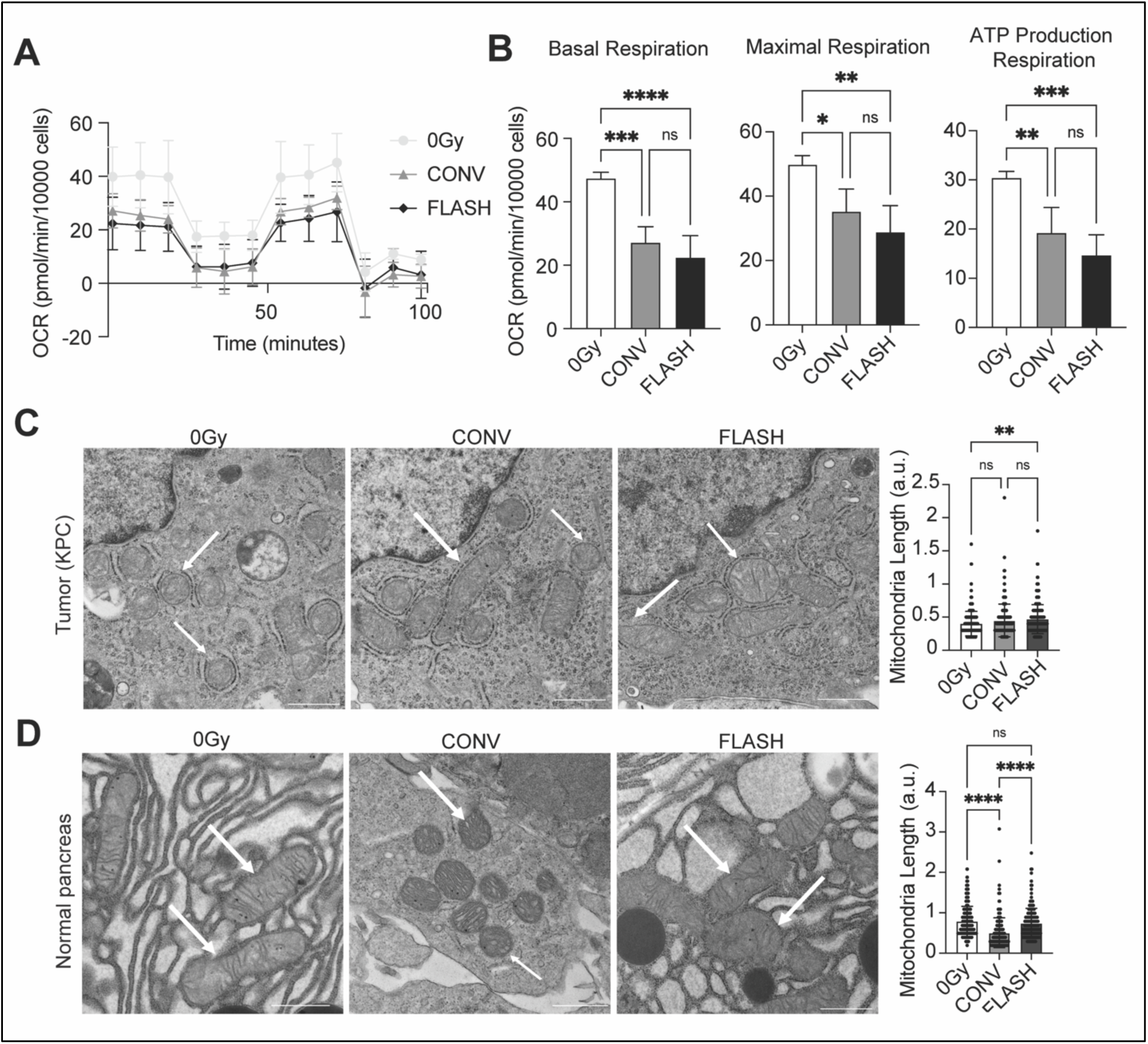
Mitochondrial effects in ex vivo KPC cells after CONV and FLASH. (A) OCR in subcutaneously injected KPC cells isolated 3.5 days after irradiation, with quantification of (B) basal respiration (left), maximal respiration (middle), and respiration linked to ATP production (right). (C) Transmission electron microscopy of mitochondria in ex vivo tumors and (D) normal pancreas 3.5 days after irradiation (scale bars 800nm, white arrows).

### FLASH extends survival in a subcutaneous pancreatic cancer mouse model

Finally, we evaluated if FLASH altered tumor growth or extended survival in an in vivo subcutaneous model of pancreatic cancer. After 20Gy CONV or FLASH, relative tumor growth was reduced compared with the unirradiated condition, but no differences were found between the two modalities (Fig. 6A). However, median survival of mice after 25Gy FLASH (12 days) was higher than after 20Gy CONV (7.5 days) (Fig. 6B). CONV-treated mice were euthanized early due to skin toxicity events, whereas unirradiated controls and FLASH did not experience these events and were euthanized due to tumor burden. Thus, survival rates are not true survival but based on animal protocol-specified parameters related to toxicity. These findings confirm that FLASH extends survival in an in vivo model of PDAC.

**Fig. 6.**
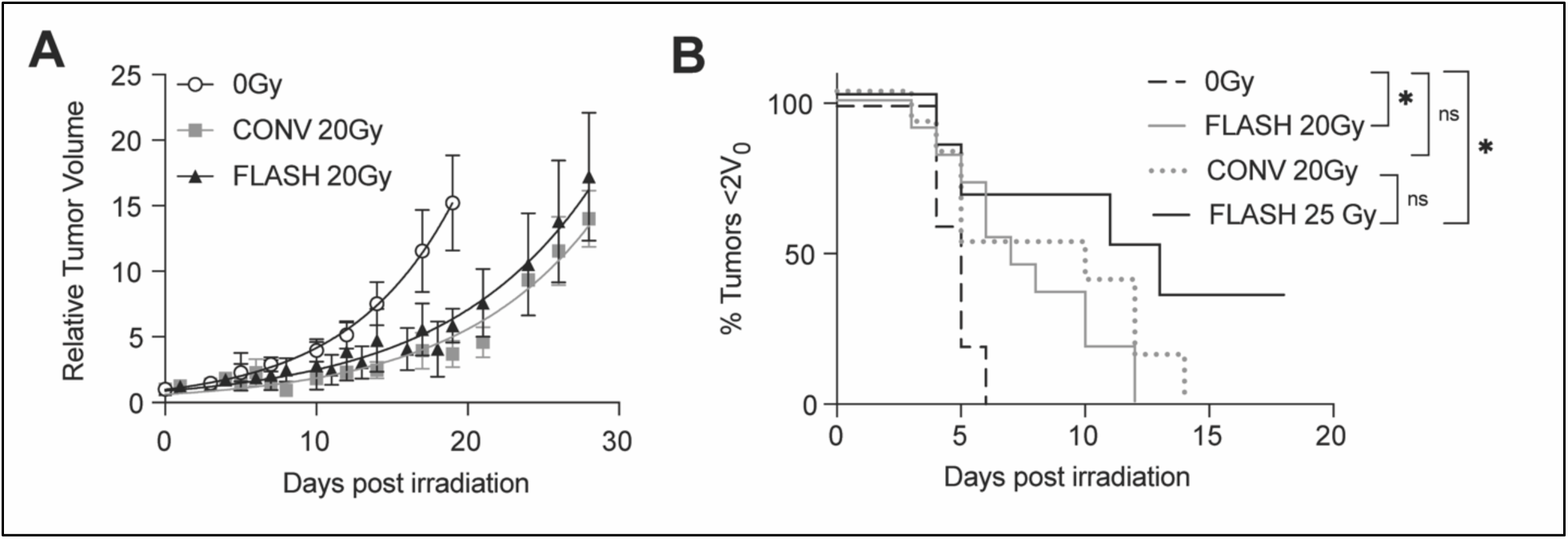
Subcutaneous model of pancreatic cancer in mice subjected to FLASH irradiation. (A) Relative tumor growth delay of KPC cells injected subcutaneously into C57BL/6 mice after 20Gy given as CONV or FLASH (unirradiated n = 5, CONV n = 10, FLASH n = 11). (B) Survival of mice with subcutaneous tumors after 20 or 25Gy of CONV or FLASH radiation.

## Discussion

To our knowledge, this is the first study to report that FLASH spares mitochondrial function and stress in normal pancreatic tissue but not in PDAC when compared to CONV. We initially explored the effects of both radiation modalities on mitochondrial biology in vitro on two different PDAC cell lines: KPC and HY24160. Although the genetic makeup of these cells is the same *(LSL-KRAS^G12D^; Trp53^fl/+^; Ptf1⍺ Cre)*, KPC is an epithelial cell line and HY24160 is a squamous cell line. Both CONV and FLASH negatively influenced several aspects of mitochondrial biology. Membrane potential decreased in KPC cells and increased in HY24160 cells, changes that indicate decreased cell function and loss of homeostasis. OCR levels decreased after RT in both cell lines, suggesting a decrease in ETC functionality. Mitochondrial superoxide levels were increased in both cell lines, indicating oxidative stress after RT. Finally, mtDNA copy number was unchanged, suggesting that the overall number of mitochondria in the cell did not change in response to irradiation. Notably, although each assay showed differences between irradiated cells and the unirradiated controls, all assays demonstrated insignificant differences between the two radiation modalities. These results are consistent with the hypothesis that FLASH evokes an isoeffective response in tumor tissue relative to CONV.

Notably, the extent and time course of mtDNA damage was different between the two cell lines. The KPC cells had largely insignificant differences in damage, whereas the HY24160 cells showed enhanced damage in mtDNA over time after FLASH versus CONV. Because mtDNA and its damage repair pathways are understudied relative to nuclear DNA damage, knowledge is limited regarding what mechanisms repair mtDNA, how it may differ between cell types, and how it is dysregulated or dysfunctional in tumors^7,35,36^. Thus, the differences in mtDNA response may reflect HY24160 being squamous and KPC epithelial, that is, one cell type may be better able to handle and repair radiation-induced mtDNA damage. Moreover, superoxide is well known to damage nuclear DNA and is generated by CONV as the main mechanism of tumor killing due to low-LET radiation. Whether mtDNA responds in the same way is unknown, but we showed that mtDNA damage occurred at the same time point that superoxide levels increased (6 hours after radiation), suggesting mtDNA may be damaged by oxygen radicals like nDNA. In comparison, nDNA was significantly damaged in KPC cells by both CONV and FLASH radiation compared to controls after 24 hours, but CONV-induced damage was significantly higher compared to FLASH. In contrast, FLASH induced significantly more damage compared to CONV in HY24160 cells. This may reflect faster damage induction with FLASH but could be followed by faster repair mechanisms. Overall, a more in-depth time course evaluation is needed to see the full extent of damage induction and repair between the two modalities. Super-resolution imaging will help accurately quantify the number of double-stranded DNA breaks at early time points after irradiation, where the expected high density of breaks will cause the intensity distributions from imaged γH2A.x to be overlapping when using conventional confocal or Airyscan microscopy^37–45^.

Finally, we explored mitochondrial morphology after RT in PDAC cells. Although irradiating KPC cells with CONV or FLASH did not change mitochondrial morphology, HY25160 cells showed significant differences in the irradiated versus unirradiated samples, and between modalities. However, KPC cells are known to have high levels of mitochondrial fission, resulting in punctate mitochondria^32^. HY24160 cells have a better balance between fission and fusion, resulting in more tubular and elongated morphology (see Fig. 4). When mitochondria are damaged or stressed, they respond with fission and eventually mitophagy, effectively budding off damaged pieces of the mitochondria that are then degraded^46,47^. This likely happened in the HY24160 cells, resulting in decreased branches per mitochondria, meaning that the cells increased fission to deal with the radiation-induced stress. However, because KPC are already in a maximal fission state^32^, it is unlikely that the mitochondria can induce more fission, resulting in the insignificant changes in morphology.

We next examined a nontumorigenic pancreatic ductal cell line, IK3025, to investigate the effects of FLASH on a cell line that cannot form tumors. We hypothesized that FLASH would lead to less mitochondrial damage than CONV in these cells. Indeed, in FLASH-irradiated cells, membrane potential, mtDNA damage, and OCR levels were maintained, as indicated by insignificant differences compared with unirradiated controls, and showed significant changes relative to CONV-treated cells. nDNA damage was significantly increased in both modalities compared to control, but FLASH caused less damage than CONV. Compared to unirradiated cells, CONV increased mitochondrial membrane potential, mtDNA damage, nDNA damage, and mtDNA copy number and decreased OCR levels, largely reiterating our findings in the pancreatic cancer cells. We conclude that FLASH largely spares these nontumorigenic cells from the damaging effect of RT, resulting in superior mitochondrial function after treatment. However, although these cells do not form tumors in vivo, they are neither primary nor truly normal cells but rather are spontaneously immortalized and harbor a p53 mutation to enable their growth in culture. Thus, although these cells reflect a FLASH effect in vitro, proper care must be taken to not overinterpret these results.

In this study, all cells were grown and assayed at normoxic (21% oxygen) conditions. Although this is standard for in vitro assays, it does not replicate the hypoxic environment that is characteristic of PDAC^48,49^. Another group found that the FLASH effect was lost in vitro under normoxia and was regained only in hypoxia^50^, but this effect was examined only in tumor cells. Because the FLASH effect is thought to be a normal tissue response, normal cells should also be examined. Another group^10^ found that in normoxia, nDNA damage was reduced in FLASH-irradiated normal fibroblasts compared with CONV-irradiated cells, demonstrating the FLASH effect in non-cancerous cells at normoxia.

To address the limitations of in vitro studies, particularly the differences in oxygen levels, we examined whether the in vitro results could be replicated in the in vivo setting. We first measured OCR and assessed mitochondrial morphology ex vivo. The OCR assay reiterated our in vitro findings, with both radiation modalities negatively affecting OCR in tumor cells relative to unirradiated controls, and no significant differences between CONV and FLASH. Thus, the in vitro normoxia findings were replicated in an in vivo physiological oxygen setting. Morphologic analysis showed that although FLASH slightly elongated mitochondria in tumor cells, it did not significantly change mitochondrial length in normal pancreas tissue compared to unirradiated tumors, whereas CONV significantly shortened mitochondria, marking a state of cell stress. We also demonstrated the FLASH effect in vivo, as relative tumor growth was unchanged, but survival rates were significantly increased after FLASH versus CONV, as CONV induced toxicity and ulceration leading to early euthanasia, which did not occur after FLASH. These findings lead us to conclude that our in vitro model, in which FLASH spared nontumorigenic cells in terms of the mitochondrial damage characteristic of CONV, can be translated to our in vivo setting, regardless of oxygen level.

## Data availability

Data will be shared upon request submitted to the corresponding authors.

## Acknowledgments and Funding

We thank Christine F. Wogan, MS, ELS, of MD Anderson’s Division of Radiation Oncology, for editorial contributions to this article. The KPC cells were a generous gift from Anirban Maitra at MD Anderson Cancer Center. The HY24160 and IK3025 cells were a generous gift from Haoqiang Ying at MD Anderson Cancer Center. This research is supported in part by the National Cancer Institute of the National Institutes of Health (NIH) under awards R01CA269565, P30CA016672, R35GM155365, and R01CA266673; by the Cancer Prevention & Research Institute of Texas (CPRIT) grants RP230224, RR200025, and RP240539, and RR200009; by the University Cancer Foundation via the Institutional Research Grant program at MD Anderson Cancer Center; by the University of Texas MD Anderson Cancer Center, Division of Radiation Oncology; by the University of Texas MD Anderson Cancer Center UTHealth Graduate School of Biomedical Sciences Dr. John J. Kopchick Fellowship; by training fellowships from UTHealth Houston Center for Clinical and Translational Sciences TL1 Program (Grants No. TL1 TR003169; T32 TR004905); and by UTHealth Innovation for Cancer Prevention Research Training Program Pre-doctoral Fellowship (Cancer Prevention and Research Institute of Texas grant RP210042). The content is solely the responsibility of the authors and does not necessarily represent the official views of the National Institutes of Health nor of the Cancer Prevention and Research Institute of Texas.

