## Supplementary Data for "Mitochondrial Responses to Conventional and Ultra-high Dose Rate (FLASH) Radiation"

**Tables S1-S5**

| Table S1. Reagents and Suppliers | | |
|---|---|---|
| **Reagent** | **Company** | **Catalog #** |
| C57BL/6 mice | The Jackson Laboratory | K8484 |
| RPMI-1640 | Corning | 10-040-CV |
| DMEM/F12 with HEPES | ThermoFisher | 11330032 |
| FBS | Sigma-Aldrich | F4135, lot 22G096 |
| Penicillin/Streptomycin | Gibco | 15140122 |
| GlutaMAX | Gibco | 35050061 |
| Sodium Pyruvate | Gibco | 11360070 |
| Insulin | Gibco | 12585-014 |
| Plasmocin prophylactic | InvivoGen | ant-mpp |
| D-(+)-glucose | Sigma-Aldrich | G7021 |
| Nicotinamide | Sigma-Aldrich | N0636 |
| 3,3’,5-Triiodo-L-thyronine | Sigma-Aldrich | T2877 |
| Dexamethasone | Sigma-Aldrich | D1756 |
| Cholera toxin | Sigma-Aldrich | C8052 |
| Insulin-transferrin-selenium | ThermoFisher | 41400045 |
| Soybean Trypsin Inhibitor | Sigma-Aldrich | T6522 |
| Mouse EGF | ThermoFisher | PMG8041 |
| Nu-Serum IV culture supplement | Corning | 355504 |
| Bovine pituitary extract | ThermoFisher | 13028014 |
| Tetramethylrhodamine | ThermoFisher | T669 |
| XF Cell Mito Stress Test Kit | Agilent | 103015-100 |
| Seahorse sensor cartridge | Agilent | 102340-100 |
| Seahorse media supplements | Agilent | 103681-100 |
| CellTak | Corning | 354240 |
| MitoSOX | ThermoFisher | M36008 |
| HBSS + Ca/Mg, no phenol red | ThermoFisher | 14025092 |
| DNeasy Blood and Tissue kit | Qiagen | 69506 |
| Hot Start LongAmp DNA Polymerase | New England Biosciences | M0534 |
| LongAmp Taq PCR kit | New England Biosciences | E5200s |
| iTaq Universal SYBR Green Supermix | Bio-Rad Laboratories | 1725124 |
| 8 well glass bottom plates | Cellvis | C8-1.5H-N |
| MitoTracker Orange CMTMRos | ThermoFisher Scientific | M7510 |
| Paraformaldehyde | Electron Microscopy Sciences | 15713 |
| Hoescht | Biotium | 40046 |
| Confocal Microscope | Leica Biosystems | LSM880 |
| MPER protein extraction reagent | ThermoFisher | 78501 |
| Bradford assay reagent | Bio-Rad Laboratories | 5000006 |
| Mini-Protean TGX Stain-Free gels | Bio-Rad Laboratories | 4568124 |
| PVDF membranes | Bio-Rad Laboratories | 1704273 |
| Blotting-grade blocker | Bio-Rad Laboratories | 170-6404 |
| SuperBlock Blocking Buffer | Fisher Scientific | 37536 |
| Clarity western ECL substrate | Bio-Rad Laboratories | 1705061 |
| Karnovsky’s fixative | Electron Microscopy Services | 15732-10 |
| Collagenase A | Roche | 1088793 |
| Hyaluronidase | Sigma | H3506 |
| Bovine serum albumin | Sigma | A9418 |
| ACK red lysis cell buffer | ThermoFisher Scientific | A1049201 |
| TrypLE | Gibco | 12605036 |
| Dispase | STEMCELL Technologies | 07913 |
| DNase1 | Sigma | 10104159001 |

| | | |
|---|---|---|
| 70μm cell strainers | Fisher Scientific | 22-363-548 |
| 40μm cell strainers | Fisher Scientific | 22-363-547 |
| Zeiss 980 Airyscan confocal microscope | Zeiss | LSM980 |

**Table S2. Beam Parameters**

| Beam Parameter | In Vitro | In Vivo |
|---|---|---|
| Dose, Gy | 4 | 20, 25 |
| No. of pulses/MU | 2 (FLASH), 500 (CONV) | 3–10 (FLASH), 1666–2923 (CONV) |
| Dose per pulse, Gy | 2 (FLASH), 0.008 (CONV) | 2.2–6.7 (FLASH), 0.008–0.012 (CONV) |
| Average dose rate, Gy/s | 360 (FLASH), 0.3 (CONV) | 265–1200 (FLASH), 0.23–0.3 (CONV) |
| Instantaneous dose rate, Gy/s | $5x10^5$ (FLASH), $6.7x10^3$ (CONV) | $1.1x10^6$–$1.6x10^6$ (FLASH), $6.4x10^3$–$1x10^4$ (CONV) |
| $T_{beam}$, s | 0.01 (FLASH), 13 (CONV) | 0.08–0.02 (FLASH), 58–97 (CONV) |
| $T_{pulse}$, μs | 4 (FLASH), 1.2 (CONV) | 2–4.1 (FLASH), 1.2 (CONV) |
| Pulse repetition frequency, Hz | 90 (FLASH), 30 (CONV) | 120 (FLASH), 30 (CONV) |

FLASH, ultra-high dose rate radiation; CONV, conventional dose rate radiation.

**Table S3. qPCR and PCR Primers**

| Gene Target | Primer Sequence (5' to 3') |
|---|---|
| 8000bp Mitochondrial DNA fragment F | ACCTGAATTGGGGGCCAACC |
| 8000bp Mitochondrial DNA fragment R | TGGCGAAGTGGGCTTTTGCT |
| mtND1 F | CAGCCGGCCCATTCGCGTTA |
| mtND1 R | GCGGAAGCGTGGATAGGATGC |
| mtND2 F | CCTCCTGGCCATCGTACTCAACT |
| mtND2 R | AGAAGTGGAATGGGGXGAGGC |
| mtND6 F | AATACCCGCAAACAAAGATCACCCAG |
| mtND6 R | TGTTGGGGTTATGTTAGAGGGAGGGA |
| 18S F | CTGAGAAACGGCTACCACATC |
| 18S R | GCCTCGAAAGAGTCCTGTATTG |

**Table S4. Thermocycler settings for qPCR and PCR**

| | mtDNA damage PCR | mtDNA damage qPCR | mtDNA copy number |
|---|---|---|---|
| Initial denaturation | 94°C, 30 seconds | 50°C, 2 minutes then 95°C,10 minutes | 95°C, 3 minutes |
| Amplification | 30 cycles: 94°C, 10 seconds; 63°C, 15 seconds; 65°C, 7 minutes | 40 cycles: 95°C, 15 seconds | 40 cycles: 95°C, 2 seconds; 60°C, 30 seconds |
| Final extension | 65°C, 10 minutes | 60°C, 60 seconds | 65°C, 30 seconds |

**Table S5. Antibodies**

| Antibody | Company | Catalog # |
|---|---|---|
| anti-Drp1 | Cell Signaling Technology | 5391 |
| anti-pDrp1-616 | Cell Signaling Technology | 4494 |
| anti-pDrp1-637 | Cell Signaling Technology | 6319 |
| anti-Erk | Cell Signaling Technology | 4695 |
| anti-pErk | Cell Signaling Technology | 4370 |
| anti-PKA | Cell Signaling Technology | 4782 |
| anti-MFN1 | Abcam | 221661 |
| anti-MFN2 | Cell Signaling Technology | 9482 |

| | | |
|---|---|---|
| anti-OPA1 | Cell Signaling Technology | 80471 |
| anti-actin | Cell Signaling Technology | 4970 |
| γH2A.x | Cell Signaling Technology | 2577 |
| Donkey anti-rabbit IgG H&L | Abcam | Ab150075 |

### Supplementary Methods

Nuclear DNA damage

KPC, IK, and HY cells were seeded in 8-well Ibidi chambers (8-well chambered glass coverslips, 80827, Ibidi) 3 days before the irradiations. The samples were then transported to an incubator near the linear accelerator (linac) and allowed to adjust for 30 minutes. The cells were irradiated to a dose of 4 Gy (CONV or FLASH), or SHAM treated before being returned to the incubator. Cells were then washed twice with 1X PBS (SH3025601, Cytiva), fixed with chilled 4% paraformaldehyde (PFA) made by diluting 16% PFA (15710, Electron Microscopy Science) in PBS for 20 minutes at 24h and 48h post-irradiation, washed once in PBS, incubated at room temperature for 10 minutes with 10 mM ammonium chloride (213330, Sigma-Aldrich) in PBS, washed once in PBS, permeabilized by washing three times with 0.2% (v/v) Triton-X (X100, Sigma-Aldrich) in PBS with 5-minute incubations, and blocked with 3% (w/v) bovine serum albumin (BSA, A2058, Sigma-Aldrich) in PBS for 1h at room temperature. The cells were then immunolabeled for γ-H2AX using primary antibodies (rabbit anti phospho-histone H2A.X (Ser139) #2577, lot 14, Cell Signaling) diluted 1:500 in 1% (w/v) BSA in PBS for 2h at room temperature, washed three times with 0.1% (v/v) Triton-X in PBS with 3-minute incubations, and then incubated with AF647-conjugated secondary antibodies (donkey anti-rabbit IgG H&L (Alexa Fluor® 647), lots: GR3312455-1 and GR3419428-1, Abcam) diluted 1:1000 in 1% BSA in PBS for 1h at room temperature. The cells were then washed five times with 0.1% (v/v) Triton-X in PBS. Finally, the cells were counterstained by incubation for 10 minutes with 300 nM DAPI (D9542, Sigma-Aldrich) in PBS and washed twice with PBS. The cells were then stored in PBS at 4°C shielded from light until imaging.

The fixed and labeled cells were imaged using a confocal microscope (Zeiss LSM980 Airyscan-2 Confocal, Plan-Apochromat 63x/1.40 NA). The imaging settings were optimized for the detection of DAPI and AF647. The following acquisition settings were used: confocal frame scanning mode, 512 pixels x 512 pixels (134.7 μm x 134.7 μm), 2.52 s frame time, and 1.02 μs pixel dwell time. DAPI was imaged in track 1 using the 405 nm excitation laser at 0.35% with an emission detection range of 411-605 nm, with a pinhole size of 1 airy unit (0.4 μm), master gain of 720 V, digital gain 1, and digital offset of 0. AF647 was imaged in track 2 using the 639 nm excitation laser at 2% with an emission detection range of 640-736 nm, with a pinhole size of 1 airy unit (0.6 μm), master gain of 780 V, digital gain of 1, and digital offset of -512. All acquisitions were averaged twice per line using mean intensity calculation with a bit depth of 16.

To quantify γ-H2AX clusters, first, manual nuclear masks were generated in ImageJ from the DAPI channel using Otsu thresholding without resetting the range. Binary Close was applied to connect nearby regions, followed by Binary Fill Holes to ensure complete nuclear segmentation. The Watershed function was then used to separate touching nuclei. Next, Analyze Particles was used with the following settings (Size: 75-500, Circularity: 0.3-1, Show: Masks, Exclude on Edges, Include Holes) to remove small, non-nuclear features in the masked images. The γ-H2AX signal was extracted from the second channel of the image, and a binary mask of γ-H2AX clusters was generated by setting an intensity threshold of 10,000 counts. Finally, the numbers of clusters per nucleus were extracted.

Mitochondrial morphology

The fixed and labeled cells were imaged using an Airyscan microscope (Zeiss LSM980 Airyscan-2 Confocal, Plan-Apochromat 63x/1.40 NA) where z stacks with 25 z positions and with a z interval of 0.16 μm were acquired. The imaging settings were optimized for the detection of Hoechst and Mitotracker Orange. The following acquisition settings were used: frame scanning with Airyscan super-resolution, 2244 pixels x 2244 pixels (79.2 μm x 79.2 μm), 1.03 s frame time, and 0.35 μs pixel dwell time. Hoechst was imaged in track 1 using the 405 nm excitation laser at 0.5% equivalent laser power in confocal mode with an emission detection range of 499-557 nm, with a pinhole size of 1 airy unit, master gain of 740 V, digital gain 1, and digital offset of 1. The filter used in the emission path was the SBS Plate 10°. Mitotracker Orange was imaged in track 2 using the 561 nm excitation laser at 0.25% equivalent laser power in confocal mode with an emission detection range of 570-620 nm, with a pinhole size of 1 airy unit, master gain of 710 V, digital gain of 1, and digital offset of 1. The filters used in the emission path were BP 420-480 + BP 570-630. All acquisitions used SR-4Y multiplexing with a bit depth of 16. Zeiss Airyscan Processing using Standard strength was applied to the images using the auto filter.
